# Pars opercularis underlies efferent predictions and successful auditory feedback processing in speech: Evidence from left-hemisphere stroke

**DOI:** 10.1101/2023.10.14.562347

**Authors:** Sara D. Beach, Ding-lan Tang, Swathi Kiran, Caroline A. Niziolek

**Affiliations:** Waisman Center, The University of Wisconsin–Madison; Department of Speech, Language & Hearing Sciences, Sargent College, Boston University; Department of Communication Sciences and Disorders, The University of Wisconsin–Madison

**Keywords:** aphasia, speech, MEG, M100, suppression, feedback

## Abstract

Hearing one’s own speech allows for acoustic self-monitoring in real time. Left-hemisphere motor planning regions are thought to give rise to efferent predictions that can be compared to true feedback in sensory cortices, resulting in neural suppression commensurate with the degree of overlap between predicted and actual sensations. Sensory prediction errors thus serve as a possible mechanism of detection of deviant speech sounds, which can then feed back into corrective action, allowing for online control of speech acoustics. The goal of this study was to assess the integrity of this detection-correction circuit in persons with aphasia (PWA) whose left-hemisphere lesions may limit their ability to control variability in speech output. We recorded magnetoencephalography (MEG) while 15 PWA and age-matched controls spoke monosyllabic words and listened to playback of their utterances. From this, we measured speaking-induced suppression of the M100 neural response and related it to lesion profiles and speech behavior. Both speaking-induced suppression and cortical sensitivity to deviance were preserved at the group level in PWA. PWA with more spared tissue in pars opercularis had greater left-hemisphere neural suppression and greater behavioral correction of acoustically deviant pronunciations, whereas sparing of superior temporal gyrus was not related to neural suppression or acoustic behavior. In turn, PWA who made greater corrections had fewer overt speech errors in the MEG task. Thus, the motor planning regions that generate the efferent prediction are integral to performing corrections when that prediction is violated.

## Introduction

Sensory feedback is central to motor control, as it delivers information about the consequences of our actions. During speech production, hearing one’s own voice allows for acoustic self-monitoring in real time. Some disorders of speech production are thought to derive, in part, from failures of monitoring and concomitantly poor auditory-motor integration (e.g., stuttering: Bradshaw et al., 2021; Cai et al., 2012; Daliri & Max, 2015; Parkinson’s disease: Railo et al., 2020; Huang et al., 2016; Mollaei et al., 2013; vocal hyperfunction: Abur et al., 2021). In post-stroke aphasia, the classic behavioral measure of auditory-motor disruption is speech repetition, where errors reflect a breakdown of the mapping from afferent speech input (the auditory target) to motor-articulatory representations (Hickok & Poeppel, 2004). The idea that the left-hemisphere lesions that cause aphasia may also damage the sensorimotor network for speech feedback processing is suggested by altered auditory feedback studies, in which persons with aphasia (PWA) make slower and/or smaller adjustments in response to externally applied pitch perturbations (Behroozmand et al., 2018; Johnson et al., 2020; Behroozmand et al., 2022). While these laboratory assays help to pinpoint the anatomical and functional bases of speech motor impairments, they fall short of addressing how PWA achieve, or fail to achieve, accuracy in their own volitional speech – in which the auditory target is speaker-internal and the feedback is veridical. As such, the current study examines the relationships among acoustic behavior, surviving neural architecture, and the cortical processing of sensory feedback resulting from one’s own speech.

Aphasic speech is often errorful, containing distorted pronunciations and perceived phonemic substitutions, omissions, and additions. Even in correct utterances, PWA tend to be more acoustically variable than their neurologically intact peers (Ryalls, 1986; Haley et al., 2001; Niziolek & Kiran, 2018). While variability alone might not be substantially detrimental to communication, there is an emerging understanding that some perceived errors stem from uncontrolled phonetic variability. That is, some phonemic paraphasias that appear to occur at the level of phonological encoding (Parsons et al., 1988) contain trace phonetic effects consistent with failures of online detection and correction of deviant acoustics during the course of articulation (Kurowski & Blumstein, 2016; also see Buckingham & Yule, 1987). These findings suggest that the “lower-order” sensorimotor network, in addition to the “higher-order” language network, may be affected in aphasia. Further, sensorimotor impairment in aphasia could be at the level of feedforward or feedback control. Analyses of acoustic variability can help to distinguish these etiologies: feedforward impairments may manifest in greater variability at syllable onset, reflecting abnormalities in top-down motor commands; feedback impairments may manifest in less online corrective action while speaking (Niziolek & Kiran, 2018). Specifically, typical speakers’ *non-errorful* speech is characterized by a phenomenon called *vowel centering*, in which productions that are initially somewhat off-target show “centering” toward the speaker’s median formant values over the course of the syllable (Niziolek et al., 2013). Moreover, there is evidence that this reduction of natural acoustic variability is reduced under masking noise (Niziolek et al., 2015, but cf. Parrell et al., 2021), highlighting the role of auditory feedback in preventing small deviations from becoming full-blown speech errors. In pilot data collected for the present study, PWA did exhibit vowel centering behavior, partially overcoming their initial feedforward variability (Niziolek & Kiran, 2018). Nevertheless, it is not well understood what surviving neural mechanisms subserve the detection and correction of deviations from intended speech sounds in PWA.

In typical brains, sensory feedback (“reafference”) is processed differently from externally-generated sensory input (“afference”). Motor cortex is thought to send an internal signal to sensory cortices, enabling prediction of the motor act’s sensory consequences. This predictive signaling has been referred to as *corollary discharge* (Sperry, 1950) and *efference copy* (von Holst & Mittelstaedt, 1950). When the efferent prediction is a good match to the true sensory input, the prediction error is small. Correspondingly, human and mammalian neurophysiology reveals that the responses of certain neural populations are suppressed during self-produced vocalization vs. passive listening to the same stimuli (Müller-Preuss & Ploog, 1981; Eliades & Wang, 2003; Creutzfeldt et al., 1989; Flinker et al., 2010; Houde et al., 2002), thus providing a general mechanism for distinguishing between self-generated (with an efferent prediction) and external (without an efferent prediction) sounds. Critical for our purposes, however, is evidence that this *speaking-induced suppression* (SIS), measured experimentally with respect to a passive-listening condition, is not all-or-none, but rather graded by the degree of match between prediction and input (e.g., Behroozmand & Larson, 2011). We have previously shown that SIS is sensitive to the “goodness” of a given utterance (Niziolek et al., 2013): That is, the neural response to off-target productions (at the periphery of a speaker’s distribution) is less suppressed than that to on-target productions (at the center of that distribution), reflecting a greater prediction error at the periphery. This fall-off of SIS from center to periphery indicates that deviant productions are *detected* by the brain. The aforementioned behavioral centering of vowel acoustics indicates that such productions are *corrected*. The purpose of the current study was to assess the integrity of this detection-correction circuit in PWA whose left-hemisphere lesions may disrupt the motor efferent prediction, the feedback comparison in auditory cortex, and/or the online control of speech.

Speech motor control is accomplished by large-scale networks across the cerebral cortex, the cerebellum, and subcortical structures, comprising feedforward and feedback control systems (Tremblay et al., 2016; Simonyan et al., 2016; Perkell, 2012). All participants with aphasia in this study had damage from a stroke in the left middle cerebral artery, but their diverse lesion patterns within its territory (Tatu et al., 1998) permitted us to relate lesion profiles to subcomponents of speech motor control. We adopted an anatomical region-of-interest approach to interrogate a putative origin and target of the efferent prediction: the left inferior frontal gyrus, pars opercularis (IFO) and the left superior temporal gyrus (STG), respectively (e.g., Rauschecker & Scott, 2009; Hickok, 2012). IFO is one of several adjacent motor planning regions, upstream of primary motor cortex, implicated in speech articulation (Blank et al., 2002; Fridriksson et al., 2016). Various instantiations of the DIVA model of speech production have placed the speech sound map, representing the sounds that speakers intend to produce, in IFO and/or adjacent ventral premotor cortex (Miller & Guenther, 2021; Guenther & Vladusich, 2012; Tourville & Guenther, 2011; Guenther, 2006). Activity in IFO precedes the suppression of sensory-evoked responses (Wang et al., 2014); therefore, IFO may be functionally analogous to the murine premotor area that has been shown to suppress ipsilateral auditory cortical activity (Schneider et al., 2014; for a review of nonhuman primate work in this area, see Eliades & Wang, 2019). STG, in addition to housing auditory cortex that performs spectrotemporal computations on speech input (Hickok & Poeppel, 2007; Bhaya-Grossman & Chang, 2022), contains the neural populations that respond to mismatches between efferent predictions and that input (Tourville et al., 2008; Niziolek & Guenther, 2013; Meekings & Scott, 2021). Moreover, overt vs. silent articulation strengthens the functional connectivity between IFO and posterior STG, perhaps reflecting the engagement of auditory feedback mechanisms for error detection and correction (Zhang et al., 2023). Left-hemisphere IFO and STG therefore served as our two regions of interest for predicting the functioning of the detection-correction circuit for volitional speech production in aphasia.

We used magnetoencephalography (MEG) to measure speaking-induced suppression (SIS) of the peak auditory cortical response, the M100, in order to characterize the efferent prediction circuit for speech and assess its integrity in post-stroke aphasia. We first compared the magnitude of SIS, an indicator of prediction delivery, among intact hemispheres (left and right in the control group, and right in PWA) and the damaged left hemispheres of PWA. We then related individual lesion profiles to SIS and to speech behavior, where vowel centering reflects feedback-guided motor control, and overt speech errors reflect failures of feedforward planning and/or online correction. Finally, taking advantage of natural, trial-to-trial variability in pronunciation, we compared the SIS of productions near the center of the vowel-formant distribution with the SIS of less typical productions at its periphery; we interpret systematic differences in SIS as an index of cortical sensitivity to the typicality of self-produced acoustics. In a secondary aim, we explored whether left-hemisphere damage was associated with atypical patterns of response lateralization across the two hemispheres. By using SIS as an objective neural measure of the efferent prediction, and considering vowel centering and speech accuracy as evidence of successful speech deviance detection and correction, we aimed to make progress on characterizing the underlying pathology of speech deficits in aphasia.

## Materials and Methods

### Participants

Fifteen persons with aphasia (PWA) and 15 age-matched controls took part in the experiment. All PWA were right-handed, monolingual speakers of American English and had chronic aphasia subsequent to a single stroke of the left middle cerebral artery. Individuals who were unable to undergo magnetic resonance imaging (MRI), had lesioned subcortical tissue, and/or had a major neurological or psychiatric diagnosis other than stroke were excluded from the study. A lesion overlap map for the aphasia group, showing the distribution of damaged voxels across the brain, is provided in **Figure 1**. PWA were 11 males and 4 females who ranged in age from 46 to 63 years. On average, they were 108 months post-stroke (SD = 111, range = 19-470). Clinical diagnoses were of Broca’s aphasia (n = 5) and anomic aphasia (n = 10). Three PWA (n = 1 Broca’s; n = 2 anomic) had a comorbid diagnosis of apraxia of speech, a disorder in which the capacity to program the movements of the articulators is impaired (Ziegler, 2008). See **Table 1** for a full characterization.

**Figure 1.**
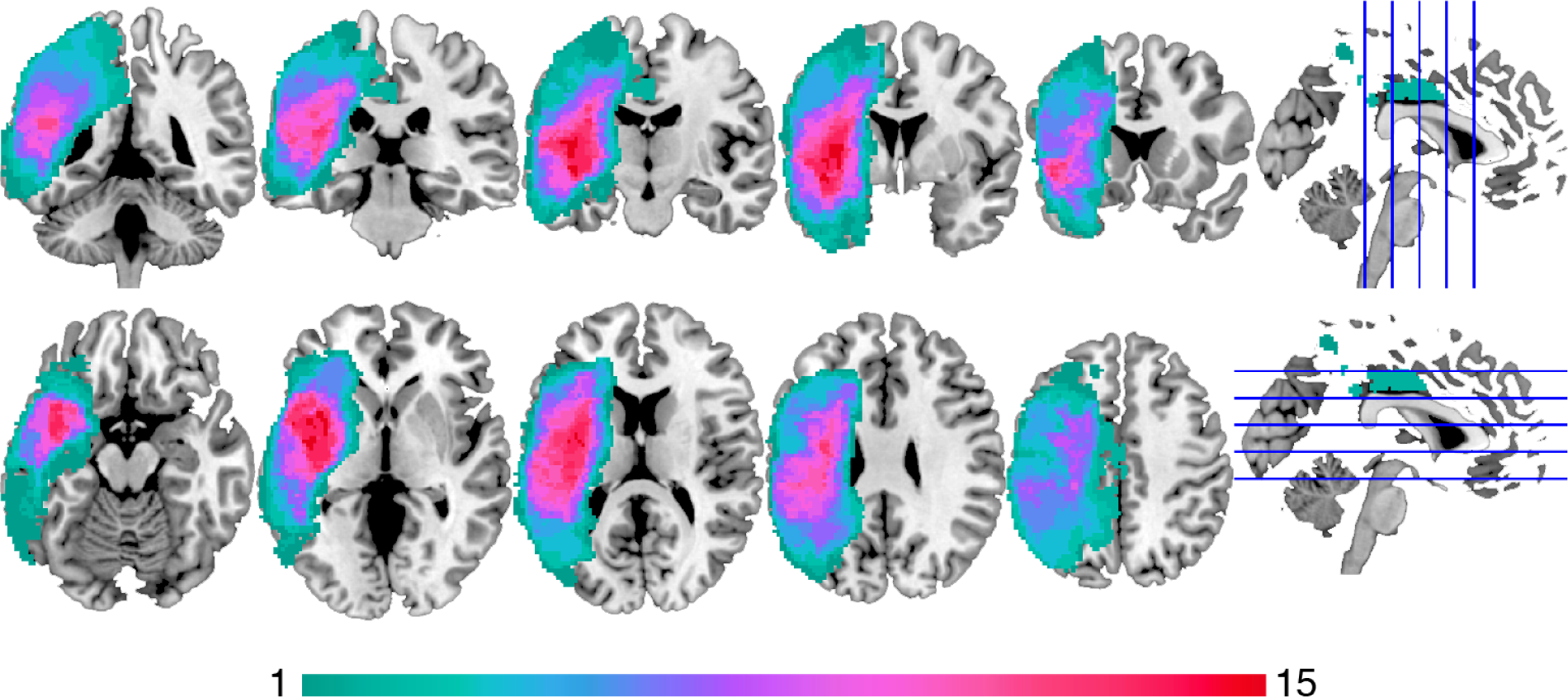
Lesion overlap map. Lesion overlap map for n = 15 PWA displayed on an MNI template brain. Warmer colors indicate a higher frequency of lesioned tissue at a given voxel.

**Table 1.** Characteristics of the aphasia group.

| PWA | age (yrs) | sex | time post-stroke (mos) | total lesion volume (mm<sup>3</sup>) | IFO spared (%) | STG spared (%) | Heschl's spared (%) | subtype | WAB AQ | PALPA7 (%) | PALPA8 (%) | PAL7 (%) | Speech errors in the MEG task |  |  |  |  |
| --- | --- | --- | --- | --- | --- | --- | --- | --- | --- | --- | --- | --- | --- | --- | --- | --- | --- |
|  |  |  |  |  |  |  |  |  |  |  |  |  | Omissions | Misselections | Distortions | Other words | Total |
| s01 | 58 | F | 49 | 56664 | 96 | 39 | 27 | anomic | 97 | 100.0 | 76.7 | 87.5 | 0 | 2 | 0 | 1 | 6 |
| s02 | 58 | M | 48 | 73912 | 100 | 51 | 7 | anomic | 82 | 95.8 | 66.7 | 70.0 | 0 | 0 | 3 | 1 | 6 |
| s03 | 52 | M | 141 | 265360 | 17 | 6 | 1 | Broca's + apraxia | 64 | 73.3 | 43.3 | 72.5 | 19 | 22 | 0 | 1 | 61 |
| s04 | 51 | M | 85 | 285800 | 28 | 16 | 5 | Broca's | 36 | 87.5 | 53.3 | 55.0 | 37 | 4 | 1 | 72 | 125 |
| s05 | 51 | M | 89 | 50464 | 95 | 94 | 81 | anomic | 95 | 95.8 | 96.7 | 95.0 | 0 | 2 | 0 | 0 | 2 |
| s06 | 57 | M | 163 | 193304 | 62 | 33 | 7 | Broca's | 50 | 91.7 | 53.3 | 65.0 | 104 | 25 | 1 | 58 | 189 |
| s07 | 54 | F | 32 | 80520 | 100 | 63 | 77 | anomic | 85 | 87.5 | 43.3 | 40.0 | 16 | 21 | 0 | 3 | 40 |
| s08 | 56 | M | 29 | 70424 | 100 | 39 | 1 | anomic | 95 | 100.0 | 66.7 | 57.5 | 0 | 2 | 0 | 0 | 2 |
| s09 | 55 | F | 117 | 217520 | 24 | 4 | 0 | anomic | 92 | 100.0 | 86.7 | 80.0 | 0 | 3 | 0 | 1 | 4 |
| s10 | 52 | M | 85 | 76792 | 75 | 29 | 12 | anomic + apraxia | 86 | 100.0 | 86.7 | 92.5 | 35 | 24 | 9 | 55 | 124 |
| s11 | 63 | M | 177 | 136464 | 42 | 45 | 21 | Broca's | 73 | 100.0 | 73.3 | 75.0 | 24 | 48 | 1 | 70 | 149 |
| s12 | 58 | F | 62 | 107872 | 14 | 89 | 71 | anomic | 85 | 83.3 | 63.3 | 72.5 | 27 | 61 | 10 | 50 | 149 |
| s13 | 46 | M | 49 | 43648 | 44 | 91 | 86 | anomic + apraxia | 94 | 62.5 | 53.3 | 50.0 | 1 | 3 | 3 | 2 | 9 |
| s14 | 53 | M | 470 | 119968 | 20 | 29 | 22 | anomic | 91 | n/a | 80.0 | n/a | 23 | 15 | 2 | 15 | 58 |
| s15 | 62 | M | 19 | 95424 | 59 | 48 | 32 | Broca's | 56 | 100.0 | 3.3 | 52.5 | 1 | 92 | 10 | 21 | 137 |

The control group was composed of 15 adults who were recruited to match each PWA on sex and age ±6 years. These individuals reported no history of speech, hearing, or language disorder and were also right-handed, native speakers of English. These 11 males and 4 females ranged in age from 43 to 69 years.

Because the study was concerned with auditory feedback processing, pure-tone audiometry air-conduction thresholds at 250, 500, 1000, 2000, and 4000 Hz were obtained to characterize hearing abilities in both PWA and controls. There were no significant group differences in thresholds in either ear at any of the frequencies tested (all *p*’s > 0.12; **Figure S1**). Two individuals in each group self-reported tinnitus.

Participants provided written and verbal informed consent to all study procedures, which were approved and overseen by the Institutional Review Boards of Boston University’s Charles River Campus and the Massachusetts Institute of Technology.

### Procedure

The study consisted of two visits for each PWA and one visit for each control participant. At the first visit, PWA completed standardized assessments and performed a behavioral-only version of the MEG speaking task to ensure that they could read and articulate the stimulus words. Some of these feasibility data have been previously reported (Niziolek & Kiran, 2018). The neuroimaging visit, scheduled on a separate day, included MEG acquisition followed by MRI scanning. All acoustic data reported here are from the speaking task performed in the MEG scanner during the second visit, as described below.

#### Standardized assessments

The speech and language abilities of PWA were assessed using the Western Aphasia Battery (WAB-R; Kertesz, 2007), the Psycholinguistic Assessments of Language Processing in Aphasia (PALPA7 word repetition and PALPA8 nonword repetition; Kay et al., 1992), and the Psycholinguistic Assessments of Language (PAL7 word and nonword repetition; Waters & Caplan, 1995). The Aphasia Quotient of the WAB-R, a 0-to-100 measure in which lower scores indicate greater global severity, ranged in this group from 36 to 97 (**Table 1**); the lowest five scores were from the individuals with Broca’s aphasia and the highest ten scores were from the individuals with anomic aphasia. All PWA made errors in the repetition of words and/or nonwords during standardized assessments (**Table 1**), making them a suitable population in which to test hypotheses about the neural bases of speech error detection and correction.

#### MEG speaking task

Participants read aloud the words “eat”, “Ed”, and “add” (chosen to elicit the vowels /i/, /ɛ/, and /æ/ with minimal jaw movement) while hearing their own speech played back in real time (*speak condition*). Words were projected on the screen one at a time in white on a black background for 1000 ms with a random inter-trial interval between 750 and 1500 ms. Participants were instructed to speak clearly while minimizing head and jaw movement. Fifty randomized tokens of each of the three words were presented, with a short break provided after every 30 trials. After the speak block, participants saw the same stimuli in the same order with the same timing and heard their prior productions played back to them (*listen condition*). Thus, the listen condition provided the same visual and air-conducted acoustic stimulation as the speak condition, absent the motor activity. There were four runs of the experiment, for a total of 600 speak trials and 600 listen trials (200 for each of “eat”, “Ed”, and “add”).

In a few cases, the paradigm was adjusted to the abilities of PWA as determined in the first visit. Participant s03 was given a longer inter-trial interval in order to minimize apparent fatigue. Participant s04 had difficulty reading the words when they were randomized and instead saw the same word presented on every trial within a block. Participant s06 had difficulty reading “Ed” and so saw only “eat” and “add” during the experiment, completing 450 speak and 450 listen trials.

#### MEG acquisition

MEG was acquired during the task on an Elekta Triux scanner located in a quiet, magnetically shielded room. Whole-head coverage was obtained with 306 channels (102 magnetometers and 204 planar gradiometers). Signals were recorded at a sampling rate of 1000 Hz and filtered online between 0.03 and 330 Hz. Continuous head position measurements were collected from five coils affixed to the scalp. These coil locations, in addition to three anatomical landmarks (nasion, left and right preauricular points) and ∼100 points on the scalp and face, were digitized using a Polhemus Fastrak system (Colchester, VT) and used to co-register the MEG and MRI scans.

Participants’ speech was captured at 11025 Hz on a Shure SM93 lavalier microphone modified for MEG compatibility and a Focusrite Scarlett 2i2 sound card. The audio was split, routed to both the participant and an input channel on the MEG scanner. Playback during both speak and listen conditions was delivered to the participant without delay via insert earphones (Etymotic, Oak Grove Village, IL) and matched in sound level across the two conditions. The scanner’s audio channel had the same sampling rate as the MEG signal, allowing for precise time alignment of neural recording and auditory stimulation.

#### MRI acquisition

Subsequent to the MEG session, a T1-weighted anatomical MRI scan (voxel size 1×1×1 mm; FoV 256 mm; 176 slices; TR 2530 ms) was obtained on a 3T Siemens Magnetom TrioTim scanner for use in estimating the neural sources of the MEG signals, and, in PWA, characterizing lesions.

### Analysis

#### Acoustic analysis

The acoustics of each spoken word were analyzed in order to measure trial-to-trial variability in production and real-time adjustments over the course of each syllable. First, errorful trials were discarded from analysis. A research assistant listened to a recording of each production, viewed the corresponding word stimulus, and marked it for rejection if it contained a perceptible error, including an omission (no vocal response), a misselection (e.g., saying “eat” for “add”), a severe vowel distortion, or a production of any other word outside the stimulus set (e.g., “head”, “had”) (**Table 1**). The remaining trials therefore included a range of pronunciations that were nonetheless perceived as correct by an unfamiliar listener. For these trials, trajectories of the first and second formant frequencies (F1 and F2) were tracked using wave_viewer (Niziolek & Houde, 2015), a Matlab interface to Praat software (Boersma & Weenink, 2019). The onset and offset of voicing were detected with an automated intensity threshold and, when necessary, manually corrected after visual inspection of periodicity in the waveform and spectrogram. Formants during the voiced syllable were estimated with linear predictive coding (LPC). The LPC order and pre-emphasis were selected to achieve stable tracking on a per-participant and per-vowel basis. Implausible values or discontinuities in the trajectories were corrected by adjusting these two values on a per-trial basis. Formants were calculated in Hz and converted to the logarithmic mel scale to accord with human perceptual sensitivity.

Time windows for formant averaging were defined as in prior studies (Niziolek & Kiran, 2018, Niziolek et al., 2013). The initial time window, from 0 (sound onset) to 50 ms, reflected the feedforward speech motor command for vowel onset. The mid-utterance time window consisted of the middle 50% of each formant trajectory, reflecting the vowel’s steady state – with sufficient time for the incorporation of auditory feedback into ongoing production. Previous studies have reported latencies of online compensation for real-time formant perturbations on the order of 150 to 160 ms (Cai et al., 2012; Caudrelier & Rochet-Capellan, 2019); adjustments for externally applied speech perturbation and speaker-internal variability are thought to rely on similar preconscious auditory feedback processing mechanisms (Munhall et al., 2009; Niziolek & Guenther, 2013; Niziolek et al., 2013).

We created trial subsets using formant values in the initial time window (0-50 ms) such that this acoustic information would precede, and therefore could contribute to, the M100 neural response. Specifically, we defined “center” trials and “peripheral” trials with respect to each participant’s median F1 and F2 values for each vowel (i.e., the center of its distribution). For each production of a given vowel, we calculated the Euclidean distance to the median in two-dimensional F1-F2 space; ***center trials*** were the n = 125 with the smallest initial distances and ***peripheral trials*** were the n = 125 with the largest initial distances (see also *MEG analysis*, below). Two key acoustic metrics, variability and centering, were defined with respect to this median. ***Variability*** was calculated as the average Euclidean distance, in mels, to the median in each time window, such that a greater distance indicates a broader distribution of formants across repeated productions of the same vowel. To determine whether speakers correct trials that are initially off-target, we calculated the ***centering*** of peripheral trials, subtracting their mid-utterance variability from their initial variability, representing the reduction in variability, in mels, from the beginning to the middle of the vowel (**Figure 2B**).

**Figure 2.**
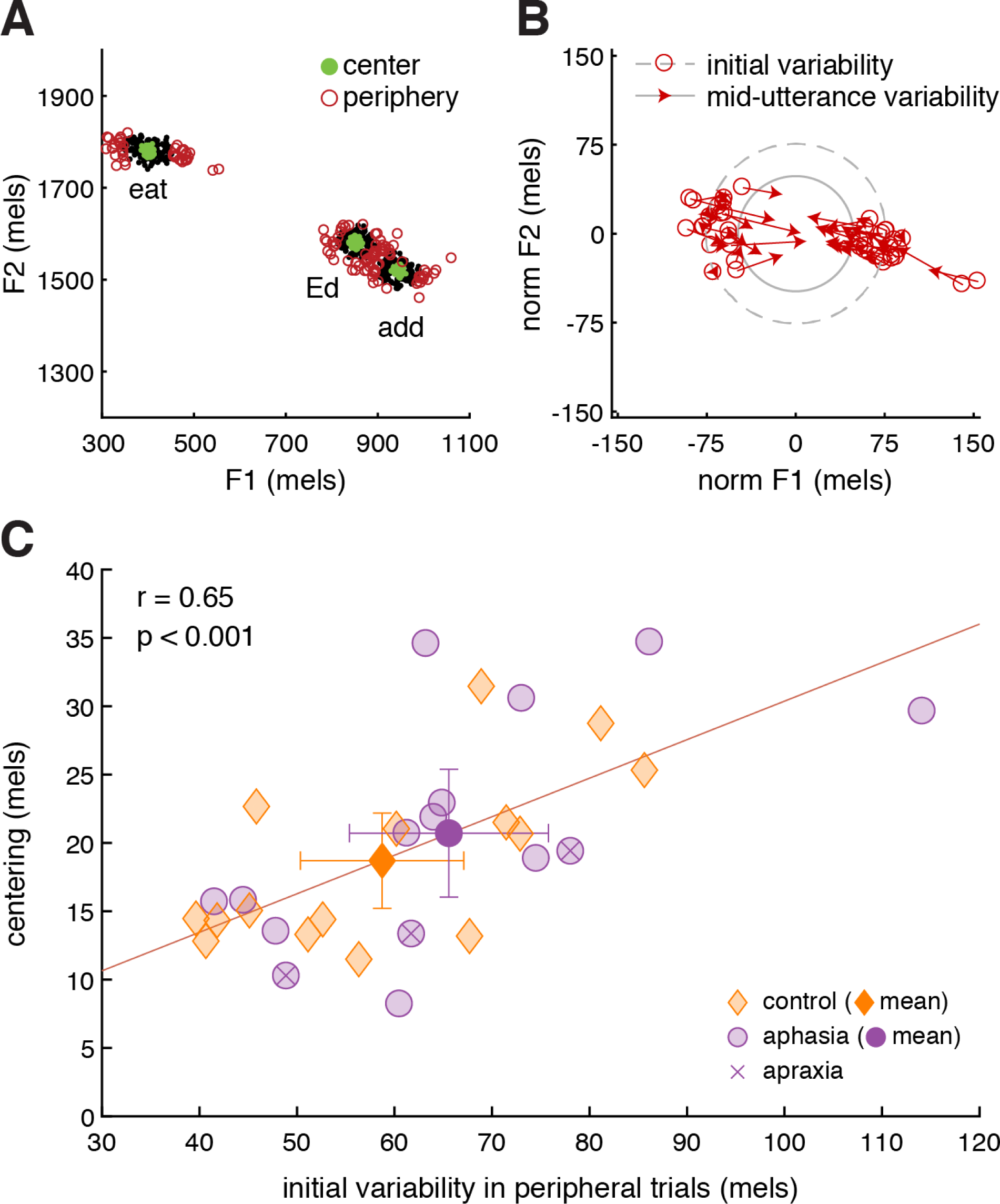
Formant variability and vowel centering. **(A)** Formant variability at vowel onset across repeated utterances of the stimulus words *eat*, *Ed*, and *add* by a representative PWA (s09). Green dots indicate center trials (n = 125 closest to the median vowel) and red circles indicate peripheral trials (n = 125 farthest from the median vowel). **(B)** Centering of the vowel /i/ in peripheral trials by a representative PWA (s09).

To ensure that centering was not simply due to random measurement error, we performed the same verification as in Niziolek and Kiran (2018): comparing the movement over time of peripheral trials (as defined above) with the movement of a different subset of trials defined as peripheral when the order of the two time windows was reversed. Specifically, this latter subset was identified as the trials with the largest *mid-utterance* distances, and we calculated their change in distance from mid-utterance *backwards* to the initial window. If centering merely reflects regression to the mean, then we would expect the movement of extreme trials to be statistically equivalent whether measured forwards or backwards in time, as indicated by a non-significant effect of time direction in an ANOVA comparing centering with time-reversed centering.

#### MEG preprocessing

Raw data were processed with Maxfilter software (Elekta, Stockholm) to remove external noise and correct for head movement. Subsequent processing was conducted in Brainstorm (Tadel et al., 2011). MEG sensors were co-registered to the participant’s MRI using six anatomical landmarks and digitized head points. Artifactual time segments were marked by hand and rejected from the continuous data. Eye-blink and cardiac artifacts were removed using signal-space projection. To reduce the influence of speaking-related movement artifacts, the data were band-pass filtered between 4 and 40 Hz. In some participants, low-frequency artifacts consistent with jaw opening and closing were evident in the sensor data even after filtering. In these cases, the topography of the artifact was used as an additional signal-space projector.

Sound onsets were automatically detected from the amplitude of the audio channel; onsets that immediately followed a visual-stimulus trigger were defined as speak and listen trial events (time = 0 at sound onset). The continuous data were epoched with respect to sound onset from −700 to 400 ms and baseline corrected between −700 and −400 ms to avoid contaminating the baseline with preparatory movement. Trials without an exact match between speak and listen conditions were excluded from analysis: For example, the corresponding listen trial was excluded for every speak trial on which the participant made an error, and the corresponding speak trial was excluded for every listen trial on which there was a severe MEG artifact. Thus, speak and listen conditions were fully matched in terms of acoustics. Out of a possible 600 total acoustic stimuli (200 for each of the three words), an average of 558 remained for controls (SD = 48, range = 440-600) and 486 for PWA (SD = 86, range = 257-594). This difference was significant, *t*(28) = 2.81, *p* = 0.01.

Neural sources were estimated from the MEG sensor data using minimum-norm imaging. The noise covariance was derived from the baselines (−700 to −400 ms) of all of the participant’s usable speak and listen trials. The forward model was computed by modeling the head as a set of overlapping spheres. Current density maps were then computed across 15,000 dipoles modeled as perpendicular to the cortical surface.

#### MRI preprocessing and lesion analysis

For all participants, the cortical surface was reconstructed from the T1 structural scan with Freesurfer (Fischl, 2012), enabling the estimation of neural sources using individual anatomy in Brainstorm as described above.

For PWA, lesion volumes and the percentage of spared tissue in anatomical ROIs were obtained in a separate processing pipeline. First, brain lesions were segmented in ITK-SNAP 4.0.0 (www.itksnap.org; Yushkevich et al., 2006), using a semi-automated method in which a Random Forests classifier was trained with six tissue labels (lesion (low intensity), lesion (medium intensity), gray matter, white matter, cerebrospinal fluid, bone) placed throughout the T1 scan. The classifier took as features the intensities of each voxel and its neighbors, as well as voxels’ coordinates, in order to distinguish the lesion from all other types of “background” tissue. Consistent with previous studies (Meier et al., 2019), enlarged ventricles were excluded from the lesion maps. Segmentation was then guided by an active contour algorithm that was seeded in the approximate center of mass of the lesion and expanded into regions where the posterior probability of the lesion was greater than that of the background (Yushkevich et al., 2016).

Subsequent processing was performed in SPM12 (http://www.fil.ion.ucl.ac.uk/spm/software/spm12/). The lesion was incorporated into the segmentation procedure by defining it as a tissue type in SPM’s tissue probability map. Warping was performed to the ICBM European brain template. The structural scan and the lesion map were then each normalized from native to MNI space, with the former using fourth-degree B-spline interpolation and the latter using nearest-neighbor interpolation to preserve its binary format.

#### Anatomical regions of interest

Anatomical regions of interest (ROIs) were defined using the AAL atlas (Tzourio-Mazoyer et al., 2002) with the MarsBAR toolbox (Brett et al., 2002). The AAL atlas, based on a normalized brain in MNI space, has been used in numerous other studies of aphasia (Yourganov et al., 2015; Shah-Basak et al., 2020; Billot et al., 2022; Behroozmand et al., 2022). In this atlas, the superior temporal gyrus is bounded by the Sylvian fissure and the superior temporal sulcus. It excludes the transverse (Heschl’s) gyrus but includes the planum temporale. The inferior frontal gyrus is bounded by the inferior frontal sulcus and the precentral sulcus. Its pars opercularis is bounded rostrally by the anterior ascending Sylvian ramus and caudally by the precentral sulcus (Tzourio-Mazoyer et al., 2002). For a post-hoc analysis in which we attempted to explain neural amplitudes in terms of primary auditory cortical tissue and hearing ability, we additionally used Heschl’s gyrus as an ROI.

As in previous studies (Johnson et al., 2019; Sims et al., 2016), calculations were performed on normalized lesion maps and brain volumes. Overlap between the patient’s lesion map and the left-hemisphere anatomical ROI was deleted and the resulting difference volume was divided by the total volume of the ROI per the AAL atlas, yielding the ***percent spared*** of each ROI. The ***total lesion volume***, in mm^3^, was also used as a predictor of speech, language, and neural responses.

#### MEG analysis

The key neural metric was the peak amplitude of the M100 auditory evoked response. Owing to the diversity of individual profiles in this study, identification of the M100 was performed manually for each participant. First, the sensor average of all listen trials and its scalp topography was examined to identify the latency of the peak auditory response. We looked for the characteristic three-peak complex of the M50, M100 and M200 (analogous to the P1, N1, and P2 event-related potentials measured using EEG) in combination with the dipolar scalp topography of the auditory response. At the time of the M100 peak, we selected the highest-amplitude vertex on the lateral surface of the brain as the seed for a functional ROI in source space. We grew this seed to a size of 10 adjacent vertices (approximately 1 to 2 cm^2^), restricted to vertices whose amplitude was at least 33% of the maximum amplitude across the cortex at that latency. When necessary, this amplitude threshold was reduced by one percentage point until 10 adjacent above-threshold vertices could be captured. This procedure was repeated in the left and right hemispheres, producing two functional ROIs per participant. The locations of these ROIs are shown as projections on the ICBM cortical surface template (**Figure 3B**). All seeds in intact hemispheres fell within perisylvian and/or temporal cortex. In lesioned hemispheres, careful efforts were made to identify the M100 in spared perisylvian and/or temporal cortex depending on the individual’s lesion profile; therefore, once projected onto the intact template brain, the left-hemisphere functional ROIs for the PWA show less spatial clustering.

**Figure 3.**
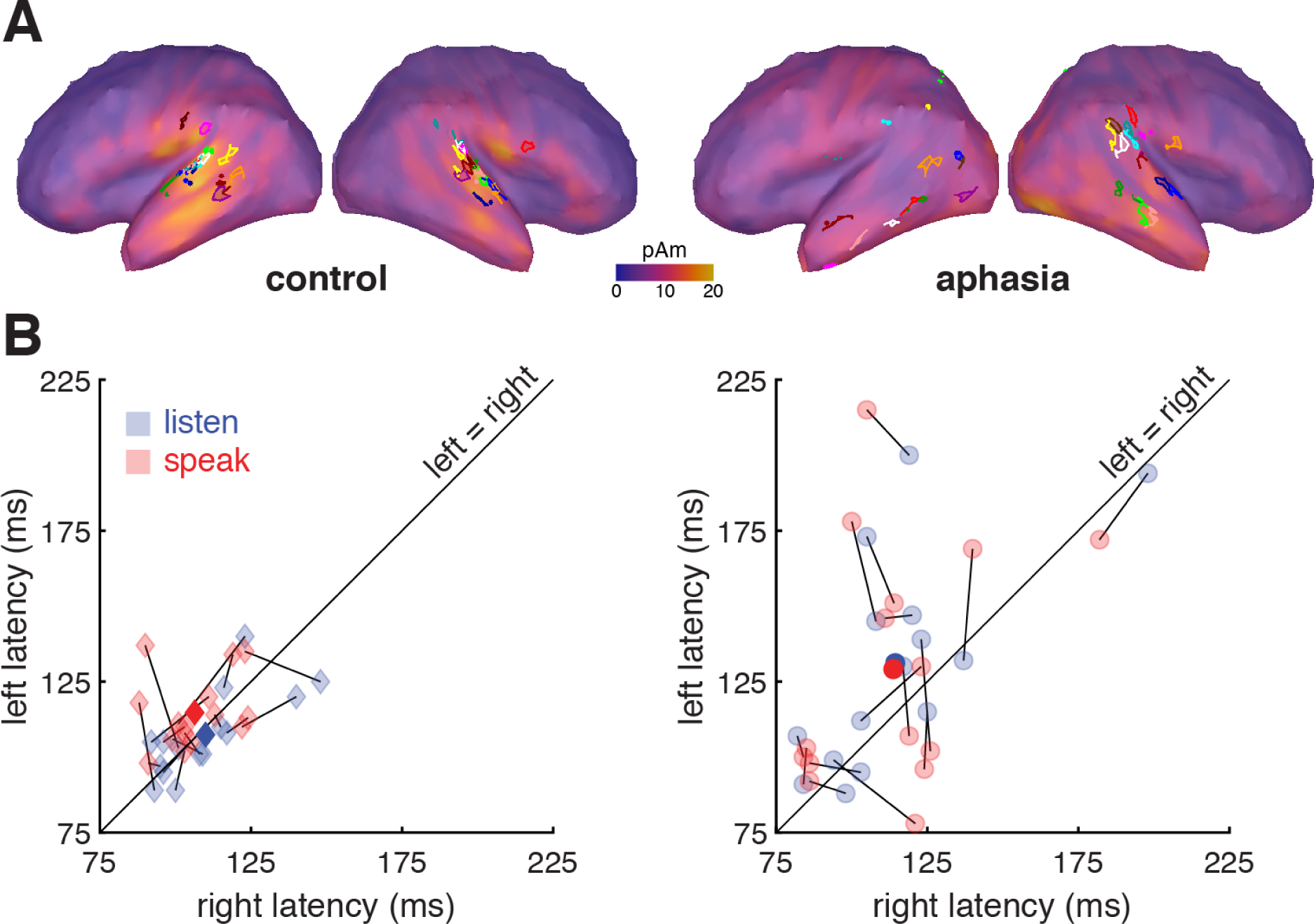
M100 neural responses identified in space and time in intact and lesioned hemispheres. **(A)** Location of functional ROIs from which M100 source space data were extracted, projected onto a common cortical-surface template. Each color represents an individual participant. In intact hemispheres, source locations cluster around canonical auditory cortex. In lesioned hemispheres, apparent dispersion of source locations reflects lesion heterogeneity across PWA as well as the unreliability of cortical projections not designed to account for lesion topology. The cortical surface shows the average listen-evoked M100 activity at the mean peak latency for that group and hemisphere. **(B)** M100 peak latencies by condition and hemisphere. Lines connect the speak (red) and listen (blue) latencies for each participant. Dark markers indicate the group mean. Data points above the diagonal indicate longer M100 latencies in the left hemisphere, which characterizes some PWA.

To obtain one time series per participant per hemisphere, we took the root mean square of the 10 vertices’ time series for the average response to all trials in the speak and listen conditions separately. Following Niziolek et al. (2013), we z-scored each time series with respect to the baseline interval of -700 to -400 ms. The M100 peak of these waveforms was then confirmed by visual inspection and established the ***peak latency*** and ***peak amplitude*** of each response. The difference in amplitude between the two conditions (listen − speak) yielded the metric of ***speaking-induced suppression (SIS)***, expressed as a z-score.

To examine the effect of acoustic typicality on neural responses, these procedures were repeated for “center” and “peripheral” subsets of speak and listen trials. Specifically, from among the trials with usable MEG data, the 125 speak trials with formants closest to the F1-F2 median, sampling the three vowels equally, were defined as the speak_center_ trials, and the 125 corresponding listen trials were defined as the listen_center_ trials. The speak_periph_ and listen_periph_ trial types were defined in the same way, using the 125 trials with formants farthest from the F1-F2 median. The previously-reported reduction in SIS for peripheral trials (listen_periph_ − speak_periph_) as compared to center trials (listen_center_ − speak_center_) is termed ***SIS fall-off*** and can be interpreted as cortical sensitivity to acoustic typicality within a vowel’s formant distribution. The number 125 was chosen so that the number of center and peripheral trials analyzed would be consistent across participants; this was limited by the participant with the fewest trials remaining after error and artifact rejection (n = 257). However, analysis of SIS fall-off was ultimately limited by the signal quality obtained during MEG recording. We could not perform the intended analysis because we could not reliably identify the M100 peak in each participant’s waveform with only 125 trials contributing to the average. Instead, we report a comparison of the center-most half and the peripheral-most half of trials.^1^

Lateralization of neural responses was evaluated using laterality indices, ranging from −1 (fully left-lateralized) to +1 (fully right-lateralized), and calculated as (right − left)/(right + left). The ***latency laterality index*** indicated the degree to which an individual’s peak latency of the listen-evoked M100 was imbalanced between their left and right hemispheres. The ***amplitude laterality index*** indicated the degree to which their listen-evoked M100 peak amplitude was imbalanced, and the ***SIS laterality index*** indicated the degree to which their magnitude of SIS was imbalanced. To prevent extreme values (Jansen et al., 2006), and consistent with our interest in neural suppression specifically, negative SIS values (i.e., speaking-induced enhancement) were replaced with 0 (“zero suppression”) for the purposes of computing laterality. It was necessary to exclude the n = 2 controls who showed enhancement in both hemispheres from the SIS laterality analysis because the denominator of the index was 0.

#### Hearing thresholds

We analyzed the contribution of hearing thresholds to neural responses evoked by speech using the ***pure-tone average (PTA)***, computed as the mean of each participant’s 500-, 1000-, and 2000-Hz thresholds (dB HL) obtained from the pure-tone audiometry in the left and right ears separately.

#### Statistical analysis

Neural responses were tested via type-III sum-of-squares ANOVA. Peak latencies and peak amplitudes of the M100 were analyzed with fixed factors of group (control, aphasia), hemisphere (left, right), and condition (speak, listen) and with a random factor of subject nested within group. The magnitude of suppression (SIS) across all experimental trials was analyzed with fixed factors of group and hemisphere and with a random factor of subject nested within group. In order to determine the presence of SIS fall-off from center to periphery, we analyzed each hemisphere separately, including only the participants who had shown SIS in that hemisphere (as these were the only participants in which we could evaluate our hypothesis) across all experimental trials. Therefore, the left-hemisphere test included n = 11 controls and n = 14 PWA, and the right-hemisphere test included n = 11 controls and n = 13 PWA. These two ANOVAs had fixed factors of group and trial type (center, periphery) and a random factor of subject nested within group. For completeness, this analysis was followed up with an ANOVA including all participants (even those without SIS), with fixed factors of group, hemisphere, and trial type and a random factor of subject nested within group. For each ANOVA described above, the significance level was 0.05 and the full model (all main-effect and interaction terms) was specified. The effect of subject was significant in all models; these random effects were omitted from the results for brevity.

Associations among variables were computed with Spearman correlation when the relationship involved one or more bounded variables (e.g., percent of tissue spared in an anatomical ROI). In all other cases, Pearson correlation was used. We assessed whether the control and aphasia groups differed on measures of variability, centering, hearing thresholds, and laterality indices by conducting two-tailed, two-sample *t*-tests. All reported *p*-values are uncorrected.

## Results

### PWA and controls center their peripheral vowels to reduce variability

We measured formant variability across all correct productions of “eat”, “Ed”, and “add” during the MEG experiment (e.g., **Figure 2A**) and found that, in contrast with prior findings (Niziolek & Kiran, 2018), PWA did not have greater initial variability than controls, neither for all trials in the experiment (*t*(28) = -1.59, *p* = 0.12) nor for the peripheral subset of trials (*t*(28) = -1.11, *p* = 0.28).

Every participant in the study demonstrated centering of peripheral trials, decreasing their average deviance over the course of the syllable (e.g., **Figure 2B**). Across participants, peripheral trials began, on average, 62 mels from the median vowel, a distance which dropped to 42 mels at mid-utterance. As in Niziolek & Kiran (2018), we confirmed that the movement of these peripheral trials over time was greater than would be expected by chance (main effect of time direction (see *Methods*): *F*(1,29) = 34.76, *p* < 0.01). The magnitude of this centering did not differ between control and aphasia groups (*t*(28) = -0.74, *p* = 0.47; M = 20 mels, SD = 7, range = 8-35). On average, centering reduced the initial dispersion of peripheral productions by 32% (SD = 9%, range = 14-55%).

In line with prior reports (Niziolek et al., 2013; Niziolek & Kiran, 2018), the magnitude of centering was positively correlated with the initial variability of peripheral trials (all participants: *r* = 0.65, *p* < 0.001; **Figure 2C**). In other words, the greater the initial spread of formant values, the greater the corrective movement. This relationship held in the control and aphasia groups separately, as well (*r* = 0.67, *p* = 0.01 and *r* = 0.62, *p* = 0.01, respectively). In **Figure 2C**, note that the n = 3 PWA with apraxia of speech fall in the middle of the group in terms of initial variability, but are relatively low in terms of centering behavior.

Formants tend to move towards the median F1 and F2 values, here normalized to (0,0), from the first 50 ms (open red circles) to the middle 50% of the utterance (black dots), thereby reducing variability. The circle radii represent the mean distance to the median in these two time windows (dashed: first 50 ms; solid: middle 50%). **(C)** Correlation between initial variability and centering. Participants with greater initial variability in peripheral trials made greater within-vowel corrective movements in those trials. The least-squares line is computed across all participants. Orange diamonds indicate control participants and purple circles indicate PWA. Error bars represent 95% confidence intervals.

### Latencies of neural responses to speaking and listening

In order to evaluate the modulatory effect of an efferent prediction on auditory cortex, we first identified the peak cortical response to self-produced speech in the speak and listen conditions. An M100 evoked response was identified in the left and right hemispheres of each participant, both control and PWA. In **Figure 3A**, the locations of these individual peak responses (functional ROIs) are shown projected onto template brains showing the mean auditory-evoked cortical activity for each group and hemisphere, captured at the mean peak latency for that group and hemisphere in the listen condition. Response latencies were faster overall in the right hemisphere (main effect of hemisphere: *F*(1,28) = 4.91, *p* = 0.03). By definition, the M100 should peak close to 100 ms after sound onset in each hemisphere, but there was a hemisphere × subject interaction (*F*(1,28) = 4.48, *p* < 0.01), indicating that the effect of hemisphere on M100 latency differed among participants. As **Figure 3B** shows, this was driven by substantially longer latencies (above the unity line) in the lesioned left hemispheres of some PWA. Overall, however, latencies did not significantly differ by group (*F*(1,28) = 2.67, *p* = 0.11). The latencies of listen- and speak-evoked responses were not significantly different (*F*(1,28) = 0.02, *p* = 0.90), and there was no hemisphere × condition interaction (*F*(1,28) = 1.41, *p* = 0.25).

### Speaking-induced suppression (SIS) of M100 amplitude

Suppression of the M100 peak amplitude in the speak condition relative to the listen condition was the primary neural metric in this study. SIS was identified to a similar degree in both control and aphasia groups and in both left and right hemispheres (main effect of condition: *F*(1,28) = 15.3, *p* < 0.01; **Figures 4A-4D**). Overall, amplitudes were lower in PWA (main effect of group: *F*(1,28) = 6.77, *p* = 0.01), but surprisingly, this was not specific to the lesioned hemisphere (see also **Figure 3A**). The effects of hemisphere and condition on amplitude varied across participants (*F*(1,28) = 5.98, *p* < 0.01 and *F(*1,28) = 7.5, *p* < 0.01, respectively).

**Figure 4.**
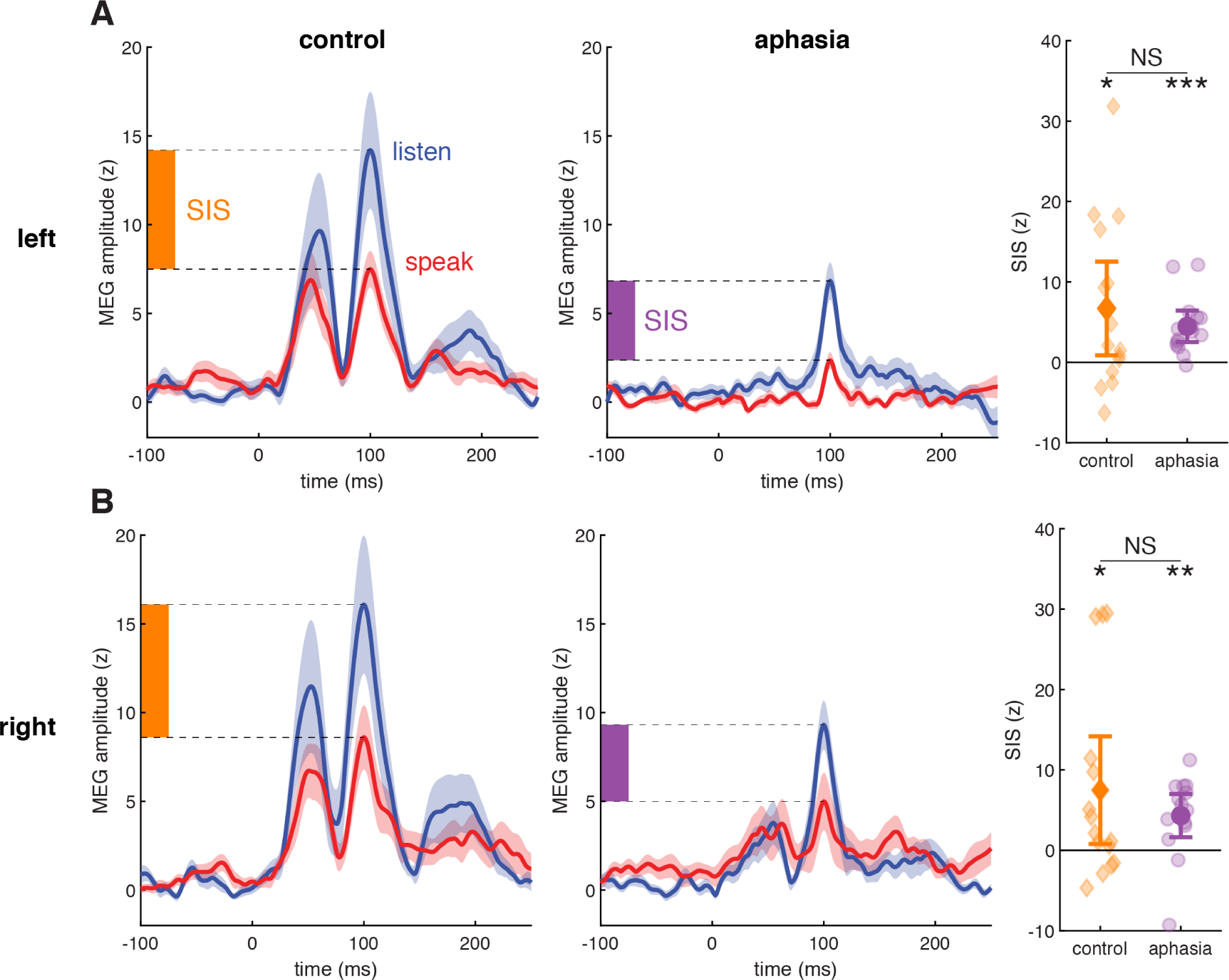
Speaking-induced suppression (SIS). **(A)** Group mean ± standard error of left-hemisphere MEG activity in listen (blue) and speak (red) conditions. Owing to the variation in M100 peak latencies across participants, individual traces have been aligned to acoustic onset (t = 0) and linearly scaled so that the M100 peak falls at t = 100. The magnitude of SIS does not significantly differ between control and aphasia. Error bars on the group-mean markers (dark orange, dark purple) represent 95% confidence intervals. **(B)** As in (A), right hemisphere. *Statistical significance*: At far right, SIS in each group and hemisphere was evaluated with a one-sample *t*-test: * *p* < 0.05; ** *p* < 0.01; *** *p* < 0.001.

The magnitude of SIS, calculated as the difference between listen- and speak-evoked peak amplitudes, did not differ by group (*F*(1,28) = 0.84, *p* = 0.37; **Figures 4E and 4F**) or by hemisphere (*F*(1,28) = 0.08, *p =* 0.78), nor was there an interaction (*F*(1,28) = 0.19, *p* = 0.66). The lack of a group effect on SIS was evident even when we excluded individuals who showed amplitude enhancement (SIS < 0) rather than suppression in the speak condition (in both hemispheres: n = 2 control; in the left only: n = 2 control, n = 1 PWA; in the right only: n = 2 control, n = 2 PWA): with these exclusions, SIS trended larger but was not significantly different in control vs. aphasia (left hemisphere: *F*(1,23) = 3.75, *p* = 0.07; right hemisphere: *F*(1,22) = 2.47, *p* = 0.13).

### Spared left IFO predicts left SIS

We took advantage of the variation in lesion profiles among PWA to ask whether the sparing of the left inferior frontal gyrus, pars opercularis (IFO) and/or the left superior temporal gyrus (STG), representing a putative source and target, respectively, of efferent suppression, was related to ipsilateral SIS. Because the sparing of an anatomical ROI predictor could be confounded with a smaller lesion overall (and therefore less severe aphasia: the smaller the lesion, the higher the WAB Aphasia Quotient, *r* = -0.70, *p* < 0.01), we additionally controlled for total lesion volume in these correlations.

Results indicated that the percent of IFO spared, ranging in this sample from 14% to 100%, was associated with the magnitude of SIS in the left hemisphere (*ρ* = 0.60, *p* = 0.02): the greater the percent spared, the greater the suppression (**Figure 5A**). We confirmed the specificity of the IFO-SIS relationship by controlling for total lesion volume; the correlation remained significant (partial *ρ* = 0.54, *p* = 0.0486). In contrast, the percent of STG spared, ranging in this sample from 4% to 94%, was not associated with the magnitude of left SIS (*ρ* = 0.17, *p* = 0.55; when controlling for total lesion volume: partial *ρ* = -0.10, *p* = 0.73).

**Figure 5.**
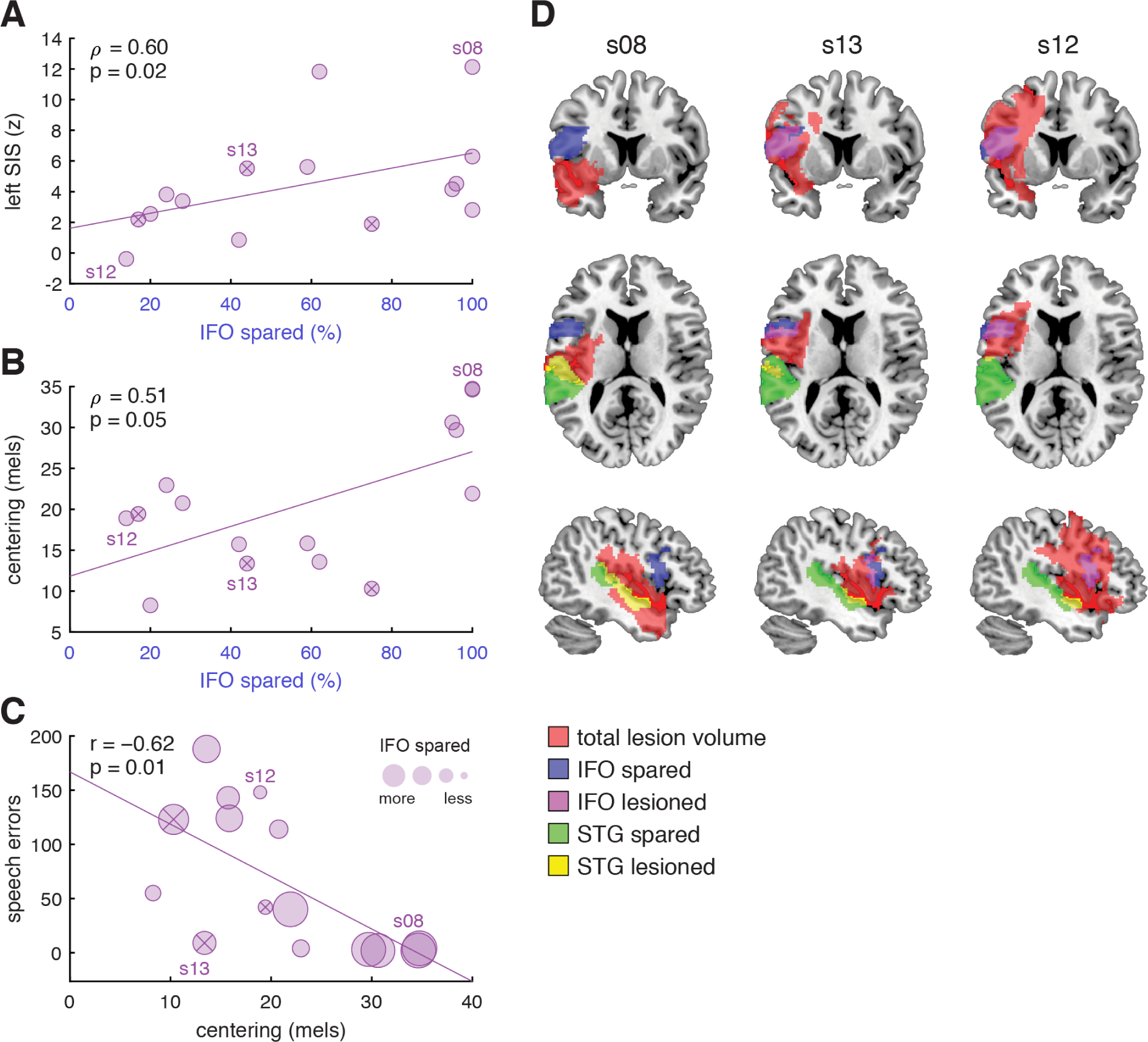
IFO predicts neural suppression and behavioral correction of vowels; correction is associated with fewer overt errors. **(A)** Correlation of percent of IFO spared and magnitude of left-hemisphere SIS in PWA. Circular markers containing an ‘×’ indicate PWA with a comorbid diagnosis of apraxia of speech. **(B)** Correlation of percent of IFO spared and magnitude of centering of peripheral trials in PWA. **(C)** Correlation of centering and number of speech errors made by PWA. **(D)** Lesion profiles of representative PWA s08, s13, and s12.

### Spared left IFO predicts vowel centering but not initial variability

We next asked whether the sparing of IFO and/or STG was related to acoustic measures of motor control, namely the formant variability measured at vowel onset and the degree to which peripheral trials were centered.

Results indicated that neither the percent of IFO spared nor the percent of STG spared were associated with initial variability, neither of peripheral trials nor of all trials (*p’s* > 0.65).

IFO spared was marginally associated with centering (*ρ* = 0.51, *p* = 0.0508): the greater the percent spared, the greater the centering (**Figure 5B**). This relationship was significant (partial *ρ* = 0.68, *p* = 0.01) when jointly controlling for total lesion volume and initial peripheral variability, which we had found to predict centering (**Figure 2B**). In contrast, STG spared was not associated with centering (*ρ* = 0.16, *p* = 0.57).

### Vowel centering (and spared left IFO) predict speech accuracy in PWA

Centering has been advanced as an objective acoustic marker of successful feedback processing, preventing deviant productions from veering across a phonetic boundary and being perceived as an error (Niziolek & Kiran, 2018). As such, we explored the relationship between centering and overt speech errors made by PWA.

We found that the greater the centering, the fewer the speech errors made during the MEG speaking task (*r* = -0.62, *p* = 0.01; **Table 2**; **Figure 5C**). This relationship remained significant when we did not include omissions (which could stem from attentional lapses or failure to initiate movement within the recording window) in the count of PWA’s speech errors (*r* = -0.57, *p* = 0.03).^2^

Furthermore, there was a strong trend such that the number of speech errors was inversely correlated with IFO integrity (*ρ* = -0.50, *p* = 0.0572), but not with STG integrity (*p* = 0.64). This can be appreciated in **Figure 5C**, where larger marker sizes indicate more spared left IFO.

Centering was not, however, associated with speech accuracy as measured on standardized assessments of repetition (PAL7, PALPA7, PALPA8, or their average; *p*’s > 0.49). Lesion profiles (IFO spared, STG spared, total lesion volume) also did not predict these standardized scores in our aphasia sample (*p*’s > 0.23).

### Representative individual data

The key findings of this study can be appreciated by considering the profiles of three representative participants with aphasia, illustrated in **Figure 5D**. In this sample of PWA, lesions affecting IFO and STG were largely dissociable: the percent spared in each region was not correlated (*ρ* = 0.37, *p* = 0.18). Two participants clearly illustrate the dissociation: s08’s lesion is relatively inferior, leaving IFO 100% spared and STG 39% spared; in contrast, s12’s lesion is relatively superior, with IFO 14% spared and STG 89% spared. In another participant, s13, IFO is 44% spared and STG is 91% spared. The left-hemisphere SIS, centering, and number of speech errors for each of these three representative participants are labeled in **Figures 5A-C**, respectively. For example, s08 has substantial left-hemisphere SIS, a large amount of centering, and few speech errors in the MEG task.

### Left SIS falls off from center to periphery of the vowel distribution

Previous research using the same paradigm in typical speakers identified greater SIS in the center vs. periphery of their formant distributions. This differential SIS was suggested to reflect neural sensitivity to natural variation in speech, such that peripheral trials showed a greater release from suppression – an effect that was specific to the left hemisphere (Niziolek et al., 2013). In our planned analyses, we intended to replicate and extend this finding by comparing SIS between center (n = 125) and peripheral (n = 125) trials in intact and lesioned hemispheres. However, we were limited by signal quality and in many participants could not reliably identify the M100 peak amongst low-level noise in the average waveforms for center and peripheral trial subsets. Instead, we took a median split and compared the magnitude of SIS in the center half of trials to that in the peripheral half, focusing on participants who, overall, showed suppression rather than enhancement in each hemisphere (left: n = 25 of 30; right: n = 24 of 30).

In the left hemisphere, there was significant SIS fall-off from the center half to the peripheral half (main effect of trial type: *F*(1,23) = 5.35, *p* = 0.03; SIS-center M = 5.70, SD = 7.25; SIS-peripheral M = 3.28, SD = 5.09). There was no interaction with nor main effect of group (*p*’s > 0.13), indicating that comparable left-hemisphere modulation by prototypicality was observed in both control and aphasia (**Figure 6**). In contrast, the corresponding analysis in the right hemisphere found no such modulation of SIS (*F*(1,22) = 0.48, *p* = 0.49). This relationship was dependent on the participants having SIS: that is, across all 30 participants, including those with enhancement rather than suppression in one or both hemispheres, an ANOVA testing SIS (with factors of group, hemisphere, trial type, and subject) revealed only a hemisphere × subject interaction (*F*(1,28) = 1.95, *p* = 0.04), no effect of group (*F*(1,28) = 2.53, *p* = 0.36), no effect of hemisphere (*F*(1,28) = 0.05, *p* = 0.86), and no significant effect of trial type (*F*(1,28) = 0.33, *p* = 0.63).

**Figure 6.**
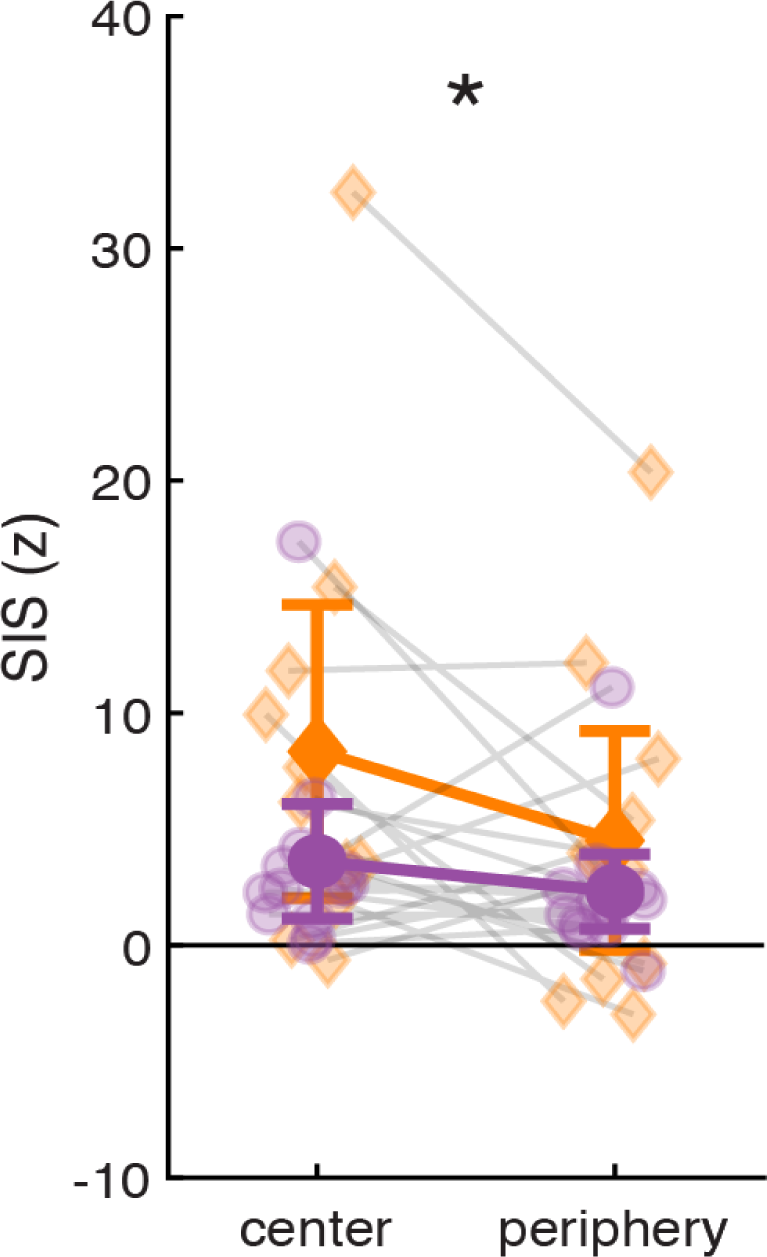
SIS is modulated by vowel typicality. Across all participants who showed suppression rather than enhancement, SIS measured in the center half of trials was greater than SIS measured in the peripheral half of trials. Error bars represent 95% confidence intervals for each group mean. *Statistical significance*: * *p* < 0.05.

Although there was no group difference in the magnitude of SIS fall-off, we investigated whether, in the aphasia group, it was related to lesion profiles. However, neither IFO spared nor STG spared were significantly associated with the magnitude of SIS fall-off, neither in the left (preserved) nor the right (potentially reorganized) hemisphere (*p*’s > 0.28).

Our final planned analysis concerned the hypothesized link between reduced SIS in peripheral trials, indicating detection of deviant acoustics via prediction error, and centering, a corrective action. Niziolek and colleagues (2013) found that, in healthy speakers, the smaller a participant’s left SIS in peripheral trials, the greater their centering behavior, suggesting that the activity in sensory regions drives the corrective action. However, this relationship was not replicated in our control sample (*r* = 0.19, *p* = 0.50). In PWA, centering was not related to peripheral SIS in either left (*r* = 0.40, *p* = 0.14) or right (*r* = 0.02, *p* = 0.94) hemispheres.

### Unexplained variation in listen-evoked amplitudes

In this study, there was notably large variation in M100 amplitudes, even though they were measured as z-scores with respect to participant-specific baseline (inter-trial) intervals, when little cognitive, perceptual, or motor activity was expected. Listen-evoked peak amplitudes ranged from 1.82 to 45.06, with the median across groups and hemispheres falling at 7.80 (M = 11.61, SD = 10.81). As the listen-evoked M100 was the reference from which SIS was measured, we attempted to account for this variation. We considered the possibility that it could be explained by individuals’ numbers of usable MEG trials (where more trials should yield a stronger neural signal) and/or hearing thresholds (where lower thresholds at speech-related frequencies (the PTA) should yield a stronger neural signal, especially given the possibility of age-related hearing loss in our participants (mean age = 56 years)). However, across all participants, the number of usable trials was not related to left or right amplitudes (*p*’s > 0.64). Moreover, across all participants, PTAs in left and right ears were not related to peak amplitudes, neither ipsilateral nor contralateral (*p*’s > 0.07).

Although PTAs did not differ between the control and aphasia groups in either ear (*p*’s > 0.2206; **Figure S1**), we paid special attention to the right-ear PTA in the aphasia group, contralateral to the lesioned hemisphere, asking whether low auditory-evoked responses here could be attributed to peripheral hearing loss and/or cortical damage. We found that the higher the PTA, the lower the listen-evoked amplitude (*r* = -0.74, *p* < 0.01 (without the outlier who had profound hearing loss in that ear; with all n = 15: *r* = -0.18, *p* = 0.52)), suggesting that peripheral hearing loss could be contributing to attenuated responses in PWA. The sparing of canonical left auditory cortex (Heschl’s gyrus and STG) was not related to the right-ear PTA (*p*’s > 0.60), so the measured thresholds likely reflected the integrity of the *peripheral* auditory system in these PWA. In fact, these auditory-cortex lesion profiles did not even predict the listen-evoked M100 amplitudes supposedly generated there (*p*’s > 0.38).

### Laterality of neural responses in PWA compared to controls

Although it was not the primary aim of our study, we took the opportunity to compute the lateralization of neural responses in the aphasia group and compare it to that obtained in an age-matched control sample. While we cannot infer a causal effect of left-hemisphere stroke on laterality given only a single, post-stroke time point, these data offer a snapshot of normative and patient populations. We explored whether lesion profiles were associated with the lateralization of responses, e.g., spared tissue supporting typical lateralization or lesioned tissue driving atypical lateralization.

#### Latency

We found that listen-evoked M100 latencies did not differ by hemisphere in controls (*t*(14) = -0.87, *p* = 0.40), but they did in PWA (*t*(14) = 2.36, *p* = 0.03); relatedly, the latency laterality index, indicating a participant’s (im)balance between the hemispheres, significantly differed between the groups (*t*(28) = 2.60, *p* = 0.01), with longer latencies measured in the left hemispheres of PWA (**Figure S2**). In PWA, the latency laterality index was associated with IFO spared (*ρ* = 0.66, *p* = 0.01) and total lesion volume (*ρ* = -0.72, *p* < 0.01), but not with STG spared or Heschl’s gyrus spared (*p*’s > 0.20). Consistent with disruption of left-hemisphere afferent pathways, more IFO spared was associated with a skew towards longer right-hemisphere latencies, and a greater total lesion volume was associated with a skew towards longer left-hemisphere latencies.

#### Amplitude

We found that listen-evoked M100 amplitudes did not differ by hemisphere, neither in controls (*t*(14) = -0.79, *p* = 0.44) nor in PWA (*t*(14) = -1.22, *p* = 0.24); relatedly, the amplitude laterality index did not differ between the groups (*t*(28) = -0.60, *p* = 0.55; **Figure S2**). Perhaps surprisingly, the amplitude laterality index in PWA was not associated with any lesion profile (IFO spared, STG spared, Heschl spared, total lesion volume; *p*’s > 0.12), so it was not that case that (right-)lateralization of evoked amplitudes was explained by left-hemisphere damage.

#### SIS

We found that SIS magnitude did not differ by hemisphere, neither in controls (*t*(14) = -0.52, *p* = 0.61) nor in PWA (*t*(14) = 0.11, *p* = 0.92). The SIS laterality index did not differ between the groups (*t*(26) = -0.32, *p* = 0.75; **Figure S2**), and thus there was no evidence that left-hemisphere stroke had altered how SIS manifested by hemisphere. The SIS laterality index (favoring the left hemisphere) was associated with IFO spared (*ρ* = -0.55, *p* = 0.03) but not with the other lesion profiles (*p*’s > 0.25).

Finally, for completeness, we noted that there were no significant correlations between any of the left-hemisphere lesion profiles (IFO spared, STG spared, Heschl spared, total lesion volume) and any of the *right-hemisphere* neural measures (latency, amplitude, SIS, SIS fall-off) (*p*’s > 0.17) that could be related to functional reorganization after stroke.

## Discussion

### Summary of results

Suppression of the auditory-evoked neural response to one’s own speech is a hallmark of feedback processing. In participants with chronic aphasia and matched controls, we found that this speaking-induced suppression was, at the group level, robust to damage to the left hemisphere. Moreover, despite overall low neural signal in the aphasia group, which we could not fully explain in terms of data quantity, hearing ability, or even lesioned tissue, we found that the magnitude of suppression in the left hemisphere was modulated by the typicality of the utterance, with less suppression measured in trials at the periphery of speakers’ vowel distributions. This finding is consistent with a framework in which the efferent prediction reflects the sensory *goal* – the prototypical vowel – such that an atypical production generates an unsuppressed prediction error.

Within the aphasia group, we found that the left inferior frontal gyrus, pars opercularis (IFO) was an important part of the feedback-processing circuit. PWA with more spared tissue in this region had greater neural suppression and greater within-syllable behavioral correction of atypical formants (“vowel centering”). Sparing of the left superior temporal gyrus (STG), conversely, was not related to either of these metrics. In turn, PWA who made greater corrective movements had fewer overt speech errors. These findings suggest that IFO is a source of the efferent signal and that its predictions underlie successful, feedback-based corrections of ongoing speech.

### IFO participates in speaking-induced suppression

The suppression of auditory reafference by frontal motor planning regions is consistent with a vast animal literature on corollary discharge (Crapse & Sommer, 2008). In particular, a direct motor-auditory connection is a structural feature of mammalian brains (Budinger & Scheich, 2009; Nelson et al., 2013; Reep et al., 1987; Frey et al., 2008), and, in the mouse, it is the secondary motor neurons that drive the suppression of auditory cortex before and during movement (Schneider et al., 2014). Human studies confirm that the source of the suppressive signal is upstream of primary motor cortex (Voss et al., 2006; Haggard & Whitford, 2004), thus linking the efferent signal to the motor plan rather than to the muscle command itself. Correspondingly, the putative plan-related signal precedes action onset in time: Source-localized EEG has revealed a correlate of the efference copy such that that activity in both left and right inferior frontal gyrus in the 300 ms prior to speaking is associated with suppression of the N1 response subsequently evoked in STG (Wang et al., 2014).

The ubiquitous phenomenon of motor-induced suppression has both general and plan-specific components. In the auditory system, general components attenuate response magnitudes within the central nervous system as well as at the auditory periphery, as early as the tympanic membrane (Carmel & Starr, 1963). Studies across mammalian species have reported auditory suppression related to general preparation and/or engagement (Li et al., 2020; Zheng et al., 2022; Otazu et al., 2009; Voss et al., 2008), and therefore, the speaking-induced suppression (SIS) measured in our study likely encompasses a non-specific suppressive phenomenon independent of forward prediction of linguistic content. However, an additional line of evidence suggests that another component of SIS is specific to the intended speech sound, above and beyond that attributed to subcortical and general-auditory dampening (Eliades & Wang, 2003; Rummell et al., 2016; Houde et al., 2002), instantiating a prediction of specific acoustic consequences. First, the fact that some neurons shift their tuning in order to monitor for deviations from expected feedback (Eliades & Wang, 2008) indicates that the vocal content, not merely the act of vocalizing, is represented neurally. Second, suppression is reduced if feedback is artificially altered to create a mismatch between predicted and actual sensation (Heinks-Maldonado et al., 2006; Behroozmand & Larson, 2011; Chang, Niziolek et al., 2013; Whitford et al., 2017). We found that PWA with more spared tissue in IFO showed greater speaking-induced suppression, suggesting that regions involved in motor planning are critically involved in sending predictions of the sensory consequences of those plans, and that this operation can be disrupted by left-hemisphere stroke.

We extend this framework – in which the specific prediction is the acoustic prototype – by likening peripheral productions to errors and showing smaller suppression in these trials. We observed this effect across controls and PWA, and in the left but not the right hemisphere, replicating the pattern identified by Niziolek and colleagues (2013) and Tang and colleagues (2023) in younger typical speakers. This phenomenon of differential SIS therefore makes a strong case that the efferent prediction is specific, suitable for comparison with sensory reafference in auditory cortex as part of the detection-correction circuit.

### IFO participates in vowel centering

We found that PWA with more spared tissue in IFO showed greater vowel centering, bringing vowel formants from the periphery of the distribution towards the median, thereby reducing variability. This relationship held even when controlling for the amount of initial formant variability across their productions. In fact, our findings suggest that damage to IFO impairs the online control of speech variability without substantially increasing feedforward variability itself.

A relationship between IFO and corrective action is consistent with prominent models of speech production that include a cortical mechanism for incorporating feedback into online control. In the State Feedback Control model, this takes the form of processing state estimate corrections after sensory error in frontal premotor areas (Houde & Nagarajan, 2011). In the DIVA model, the feedback control map, also localized to premotor planning regions, transforms error into updated articulatory trajectories (Tourville & Guenther, 2011). These models are informed by neuroimaging results in which, for example, activity in superior temporal and inferior frontal areas during formant perturbations correlates with participants’ compensatory actions (Niziolek & Guenther, 2013).

In the non-speaking mouse, homologous higher-order motor regions are recognized for their causal role in the flexible updating of motor plans subsequent to sensory feedback (Gremel & Costa, 2013; Barthas & Kwan, 2017). Causal evidence in humans is rare, but a recent study identified greater behavioral compensation and enhanced neural responses to artificially-altered pitch feedback as an after-effect of transcranial alternating current stimulation over the left inferior frontal gyrus (Li et al., 2023). Together, these lines of evidence suggest that the motor-planning regions that create the efferent prediction are integral to performing the correction when that prediction is violated.

It is important to note that several studies implicate ventral precentral gyrus more strongly than its rostral neighbor IFO in driving suppression and causing compensatory movements during speech. Connectivity analyses of human electrocorticography (ECoG) identified the ventral precentral gyrus as a source of the corollary-discharge signal, peaking prior to articulation and targeting STG (Khalilian-Gourtani et al., 2022). In that study, activity in inferior frontal gyrus preceded activity in precentral gyrus, but the stronger correlate of a motor-to-auditory discharge arose from precentral gyrus. In another ECoG study, compensation for vocal pitch perturbation was correlated with the activity of electrodes in posterior STG as well as ventral precentral gyrus, suggesting that corrective movements have premotor origins (Chang, Niziolek, et al., 2013). In that study, most electrodes in IFO did not show enhanced activity in response to speech perturbation. However, IFO rather than precentral gyrus was our *a priori* frontal ROI in the present study not only because of its role in speech motor planning (Rong et al., 2018; Papoutsi et al., 2009; Long et al., 2016; Mugler et al., 2018), but also because of its anatomical specificity. In the aphasia-standard AAL parcellation used for calculating the percent spared of ROIs, precentral gyrus contains primary motor as well as premotor cortex and is therefore not suitable for interrogating specific contributions of motor *planning* regions to the detection-correction circuit. Given the reciprocal functional connections among frontal motor- and speech-related areas (Greenlee et al., 2004), we expect that IFO and ventral precentral gyrus both participate in updating speech trajectories during online control.

Finally, it is also worth underscoring that at the group level, PWA – a majority of whom had damage to left IFO – exhibited a significant degree of vowel centering, comparable to that of controls. One potential explanation for this result is that right-hemisphere premotor regions, which were fully spared in our sample of PWA, also support corrective behavior. Whereas speech motor programs tend to be left-lateralized (Ghosh et al., 2008) and error detection tends to be bilateral (Tourville et al., 2019), there is evidence that compensation for altered feedback is related to premotor activity in the right hemisphere, especially that which is driven by right posterior temporal cortex (Tourville et al., 2008; Toyomura et al., 2007; Floegel et al., 2020; Liu et al., 2023). Findings such as these ultimately led the developers of the DIVA model to place the feedback control map in the right hemisphere (Tourville & Guenther, 2011). Our study was designed to examine the left-hemisphere circuit because of the aphasic population we were working with, but future studies could evaluate whether right-hemisphere stroke is associated with greater impairment in the online control of speech, as the DIVA model might predict.

### Vowel centering relates to speech accuracy

Vowel centering is one of a number of ways to measure speech motor control. This analysis quantifies speakers’ responses to their own natural variability and treats peripheral productions as error-like. We found that, in PWA, greater centering was associated with fewer overt speech errors, suggesting that it has functional consequences for successful production. Centering potentially prevents acoustic deviations from becoming full-blown speech errors, crossing a perceptual boundary and being perceived by listeners as a vowel misselection. This is the first demonstration of a relationship between centering, a subclinical variable, and speech accuracy, a highly salient variable. We therefore propose that centering may be a manifestation of successful feedback processing in natural speech.

Another way to assay feedback processing is to manipulate auditory feedback in real time as it is played back to the speaker. Behroozmand and colleagues (2018) found that PWA respond to external alterations of vowel pitch as controls do – by altering their true pitch to oppose the shift – but to a lesser degree. Diminished compensation was predicted by damage to left superior temporal, middle temporal, inferior frontal, and supramarginal gyri. Pars opercularis of the inferior frontal gyrus was implicated in the time window of the steepest slope of compensation, which the authors linked to the greatest need for corrective motor precision. Thus, left inferior frontal cortex has now been linked to corrective responses to both artificial pitch perturbations and naturally occuring peripheral vowels.

Like compensation for altered auditory feedback, centering theoretically relies upon the inverse model – the sensory-to-motor transformation that establishes how to move the articulators to produce the desired correction. However, there is an important distinction between the two metrics. (Indeed, in typical speakers, compensation and centering have not been shown to correlate (Niziolek & Parrell, 2021).) Altered auditory feedback paradigms (e.g., Houde & Jordan, 1998; Purcell & Munhall, 2006) manipulate auditory but not somatosensory feedback, such that speakers encounter a conflict between the two sensory modalities as well as the intended auditory prediction error. The centering analysis in the present study is used to characterize natural speech in which speakers receive veridical feedback, both auditory and somatosensory. A typical speaker might prefer to rely on one or the other (Lametti et al., 2012); a PWA might be limited by their surviving neural architecture. Whereas altered auditory feedback is very likely to drive an STG-dependent auditory error response, natural variability could be detected and corrected by means of somatosensory error. This may be one reason we did not observe a relationship between STG integrity and centering in natural speech.

Speech repetition tasks have also been used to tap the inverse model in aphasia. The ability to perceive and repeat a word, particularly a pseudoword, has been associated with larger compensation for altered feedback (Sangtian et al., 2021). We did not find a correlation between centering and standardized measures of speech repetition, which may reflect low power in our study. On the other hand, because the magnitude of centering did not differ between control and aphasia groups, it is reasonable that there would be no relationship between centering and aphasia-specific clinical assessments.

### Apraxia of speech

We paid special attention to the n = 3 PWA who had a diagnosis of apraxia of speech comorbid with aphasia because apraxia is, specifically, a motor speech disorder. These individuals did not seem to drive or disrupt the relationships we found in the larger aphasia group in terms of neural suppression, motor control, or speech accuracy. Since this population tends to have lesions in left inferior frontal gyrus (Ballard et al., 2014), we might predict low neural suppression and low centering due to IFO damage. Empirical findings regarding feedforward and feedback control systems in apraxia of speech are, to date, relatively limited and somewhat inconclusive (Maas et al., 2015; Ballard et al., 2018). Yet it is commonly said that persons with apraxia “know what they want to say and how it should sound” (Ziegler, 2008), leading them to make multiple attempts to produce intended words. This behavioral profile hints at preserved feedback-monitoring capabilities in the face of impaired motor programming. Such a dissociation – an ability to detect error but not to correct it – was not observed in the current small-scale study of a diverse aphasic population, but is theoretically possible. It also underscores the important distinction between speech errors consciously perceived – or not perceived – by the affected individual (e.g., semantic errors, phonemic paraphasias) and subphonemic “errors” in vowel pronunciation that are corrected, preconsciously, by speech motor circuitry – the focus of the present study. As proposed by Niziolek and Kiran (2018), centering analysis may complement other techniques used by speech-language pathologists to evaluate disordered speech because it reveals ongoing, feedback-guided correction that is below the perceptual threshold.

### Neural response profiles do not easily distinguish control and aphasic populations

Although PWA had lower auditory-evoked response magnitudes than controls, these low amplitudes were not confined to the damaged hemisphere, nor did they result in significantly smaller average magnitudes of suppression when comparing speak and listen conditions. This underscores a general pattern of non-significant differences at the group level, with greater variability within than across groups, and within than across hemispheres in the aphasia group. For example, we expected that listen-evoked responses would be delayed and weaker in the left vs. the right hemispheres of PWA; however, while left-hemisphere responses were slower, they were not smaller in amplitude. This bilateral pattern of responsiveness ultimately made the PWA look much like controls, whose latencies, amplitudes, and SIS magnitudes, at the group level, did not differ by hemisphere (**Figure S2**). This contrasts with previous reports, in younger, typical speakers, of greater SIS in the left vs. the right hemisphere (Curio et al., 2000; Houde et al., 2002; Niziolek et al., 2013). This heterogeneity in “typical” lateralization precludes attributing any lateralization observed in PWA to their stroke and/or recovery. For example, that the sparing of IFO predicted SIS laterality was interesting, but less so in light of having found no group difference in SIS laterality. All in all, there was very limited evidence for lesion effects on lateralization or right-hemisphere plasticity in these data.

Relatedly, lesioned tissue did not straightforwardly explain certain basic auditory response profiles in PWA. For example, the integrity of left STG and left Heschl’s gyrus did not predict hearing thresholds, confirming that tone detection survives substantial damage to these regions (Masterton & Berkley, 1974; see **Table 1**). However, the sparing of auditory cortex also showed no relationship with evoked amplitudes. While low neural signal in PWA was potentially partially explained by peripheral hearing loss, this relationship was less compelling in light of the fact that their hearing did not differ from controls’ (**Figure S1**). This unanticipated result could be addressed in future work providing a more comprehensive picture of surviving neural architecture in auditory cortex. Specifically, it would be desirable to have measures of sensory and perceptual acuity as a function of spared STG (e.g., Robson et al., 2012; Robson et al., 2013), as impairments in these domains could limit the resolution of the comparison between the efferent prediction and sensory reafference. In addition, tractography (e.g., Breier et al., 2008), functional connectivity (e.g., Schofield et al., 2012), and a finer anatomical parcellation of STG (e.g., the planum temporale: Hickok et al., 2009) would all help to elucidate the components of the cortical feedback circuit for speech in individuals with and without brain damage.

### Caveats

The significant structure-function predictive relationships within the aphasia group (IFO-SIS and IFO-centering) are tempered by the fact that although PWA, as a group, had strictly less tissue in IFO, they did not show statistically less SIS or centering than controls. Thus, one could argue that the speech error detection-correction circuit is not necessarily impaired after left-hemisphere stroke. Dysfunction relative to age-matched peers was evident in the number of errors made during a simple word-production task, but not in our subclinical acoustic measures of speech motor control (initial formant variability, a proxy for feedforward control, and centering, a proxy for feedback control), underscoring that profiles such as “high speech variability” and “low online correction ability” are not indicative of an aphasia diagnosis. On the other hand, the lack of group difference in the magnitude of SIS, and the relationship between spared IFO and SIS, is complicated by the unexpected presence of speaking-induced *enhancement* in our data (i.e., SIS < 0 in **Figure 4**). Despite overwhelming evidence across species for neural suppression during self-vocalization (Eliades & Wang, 2003; Müller-Preuss & Ploog, 1981; Creutzfeldt et al., 1989; Suga & Shimozawa, 1974), a subset of participants in our study, both controls and PWA, demonstrated larger M100 amplitudes during speaking relative to listening. (While we expected to find SIS in all controls and did not, all PWA did show SIS in one or both hemispheres.)^3^ With only a single MEG session per participant, it is unclear whether being an “enhancer” vs. a “suppresser” is a stable individual trait. Speaking-induced enhancement could potentially be attributed to individuals’ having particular cortical topography and/or a particular position in the MEG helmet that led to the recording of neural populations that are excited rather than suppressed by the act of vocalizing (Eliades & Wang, 2008). As MEG has limited spatial resolution in this regard, this is fully speculative. It is also possible that enhancement was the result of a particular cognitive or attentional state during speaking, as the M100 is obligatory but its amplitude does increase with attention (Woldorff et al., 1993). At present, it is unclear whether amplitude enhancement is functionally relevant for speaking.

### Conclusions

We interrogated sensorimotor function in aphasia, focusing on efferent predictions of the sound of one’s own speech and the detection and correction of natural variability in vowel production. Post-stroke sparing of pars opercularis of the left inferior frontal gyrus was associated with greater speaking-induced neural suppression (an indicator of efferent prediction delivery), greater vowel centering (an indicator of feedback-guided correction), and fewer speech errors. These findings highlight the importance of pars opercularis, in speaking humans, to the efferent prediction circuit that has, to date, largely been studied in non-speaking animals. Rather than initiating them, pars opercularis appears to underlie the updating of motor plans for speech; our findings suggest that it creates the efferent prediction and also performs the correction when the prediction is violated.

## Acknowledgements

We thank the staff of the Athinoula A. Martinos Imaging Center at the McGovern Institute for Brain Research (MIT) for technical support. We thank Isaac Falconer for assistance with the lesion analysis pipeline.

## Data and code availability statements

The data supporting this study are openly available at https://osf.io/hjwsx/. The code supporting this study will be made openly available upon acceptance.

## Supplementary Materials

**Figure S1.**
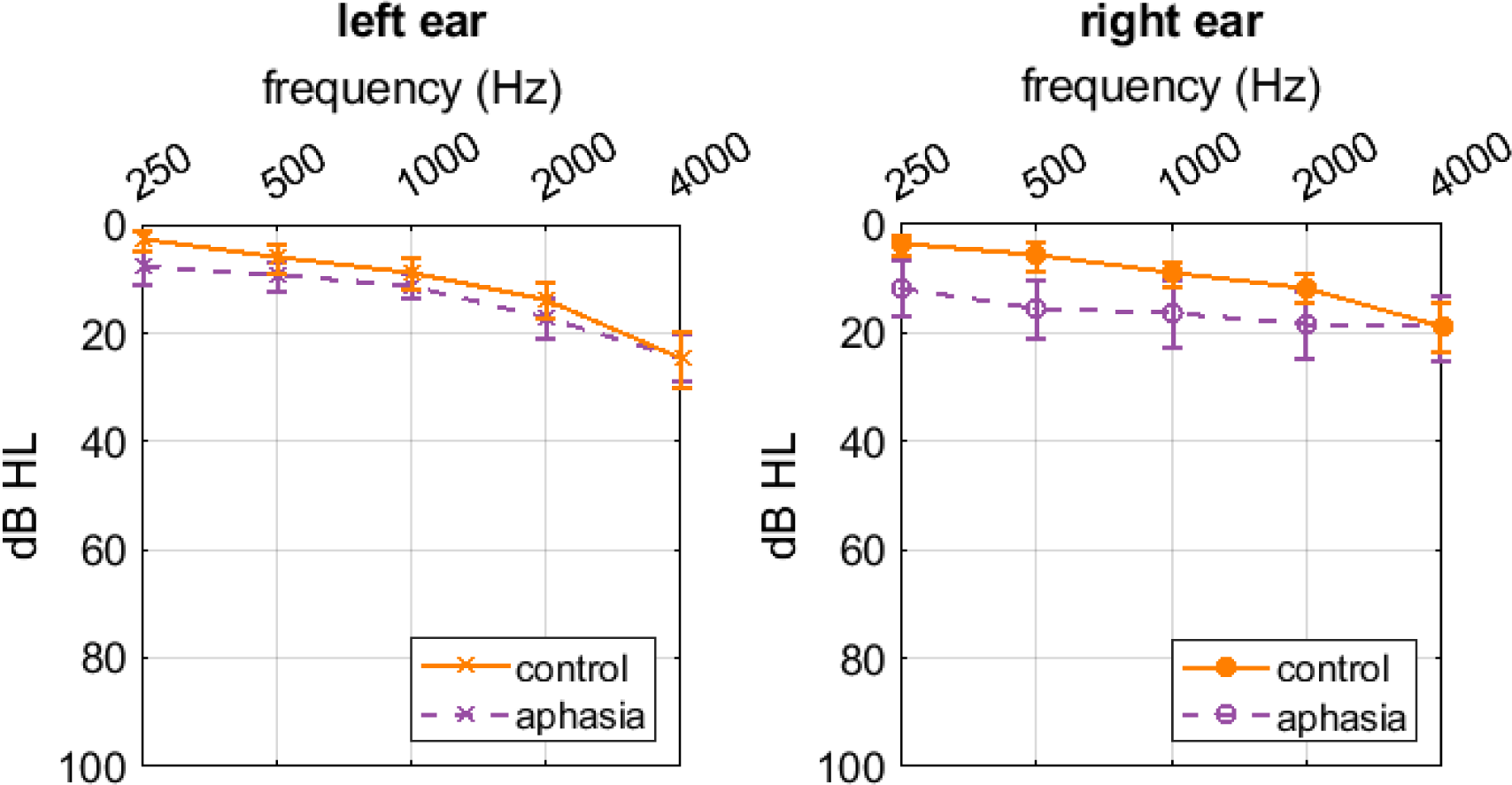
Audiograms for left and right ears. Hearing thresholds did not differ between controls and persons with aphasia at any frequency. Error bars represent the standard error.

**Figure S2.**
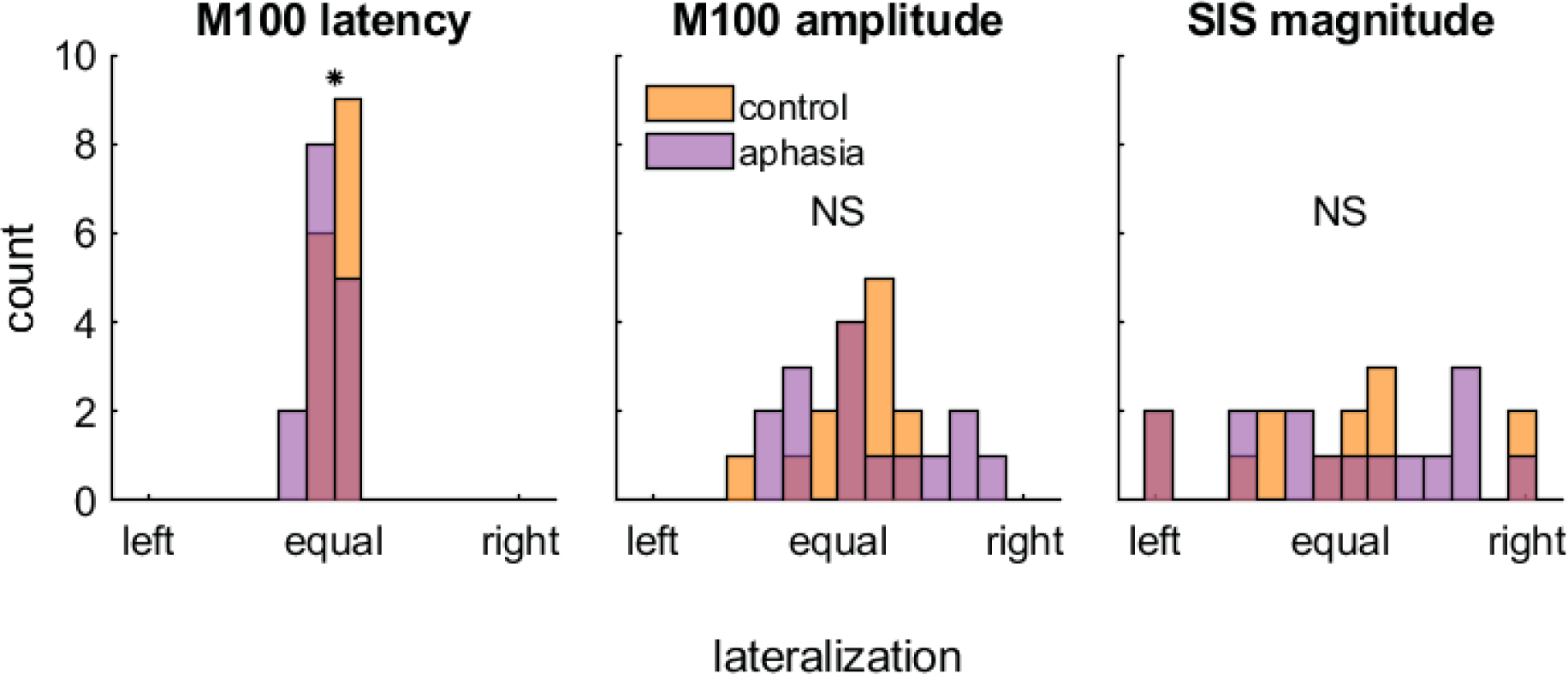
Laterality of neural responses. Each panel is a histogram depicting control (orange) and aphasic (purple) participants’ laterality index for a given neural metric. Each laterality index is calculated as (right − left/(right + left), giving a range of values between fully left-lateralized and fully right-lateralized. **(A)** The laterality of listen-evoked M100 latencies significantly differed between control and aphasia, due to longer latencies in some lesioned left hemispheres. **(B)** The laterality of listen-evoked M100 amplitudes did not differ between the groups. **(C)** The laterality of the magnitude of speaking-induced suppression (SIS) did not differ between the groups. *Statistical significance: * p* < 0.05.

## Footnotes

1 The tradeoff for more trial data is that this median split compares trials that abut one another in acoustic vowel space, decreasing the acoustic difference and therefore the hypothesized neural difference between them. And, paradoxically, including more trials potentially exacerbates the issue of data loss in PWA who made many speech errors: these individuals had fewer trials in their center half and peripheral half, making their average evoked response less reliable than that of a participant with all 600 trials available.

2 The majority of control participants made either 0 or 1 errors during the MEG task (median = 1, M = 3.73, SD = 8.37), so we did not analyze speech errors in this group.

3 Despite the heterogeneity in individual responses, the group-mean SIS effect that we measured is in line with prior reports. SIS as a percentage reduction from the listen peak to the speak peak (calculated from the grand-average, all-trials waveform in typical speakers) is 30% (Ventura et al., 2009), 39% (Houde et al., 2002), 51% (Curio et al., 2000), and 58% (Behroozmand & Larson, 2011). The corresponding value in our study (controls’ left hemisphere, **Figure 4A**) is 47%.

